# *Rhopilema nomadica* in the Mediterranean: molecular evidence for migration and ecological hypothesis regarding its proliferation

**DOI:** 10.1101/2024.09.06.610623

**Authors:** Zafrir Kuplik, Hila Dror, Karin Tamar, Alan Sutton, James Lusana, Blandina Lugendo, Dror Angel

## Abstract

Since it was first observed in Israel in the 1970s, the ‘nomad jellyfish’ *Rhopilema nomadica* has established a reputation as one of the worst invasive species in the Mediterranean Sea. It was assumed to originate in the Red Sea, or in the Indo-Pacific region, but in the absence of additional reports of live specimens outside the Mediterranean, its origins remained a mystery. Here, via molecular analysis, we present the first verified results of the existence of *R. nomadica* in the Western Indian Ocean. Moreover, using additional evidence from *Cassiopea andromeda* and *R. nomadica*, we propose that the construction of the Aswan High Dam may have led to the proliferation of *R. nomadica* in the Levantine Basin.

## Introduction

The scyphomedusa *Rhopilema nomadica* was first reported in coastal waters of the south eastern Mediterranean Sea in the 1970s (Galil et al. 1990) (Fig. 1A, 1B). By the mid-1980s, large aggregations of this species appeared annually along the Israeli coast, mainly during the summer (Spanier 1989, Lotan et al. 1992, Galil & Zenetos 2002). Thereafter, *R. nomadica* expanded north and westward into the central Mediterranean, appearing in Turkey (Öztürk & İşinibilir 2010), Greece (Siokou-Frangou 2006), Malta (Deidun et al. 2011), Tunisia (Daly Yahia et al. 2013), and more recently, Sardinia and Sicily (Balistreri et al. 2017). Dense swarms of *R. nomadica*, > 10 individuals per square meter (Angel et al. unpublished, Fig. 2), encompassing many kilometers, cause environmental, economic, and social damage, reducing biodiversity (Streftaris & Zenetos 2006), clogging water intake pipes of coastal infrastructure (Rilov & Galil 2009, Rahav et al. 2022), causing damage to the fishing industry (Nakar et al. 2011, Angel et al. 2016), and disrupting coastal recreation and tourism (Ghermandi et al. 2015). As a result of its expansion and the effects of the swarms, *R. nomadica* is considered one of the “100 worst invasive species” in the Mediterranean (Streftaris & Zenetos 2006, Zenetos et al. 2010) and one of the most “impacting species” in European seas (Katsanevakis et al. 2014).

**Fig. 1.**
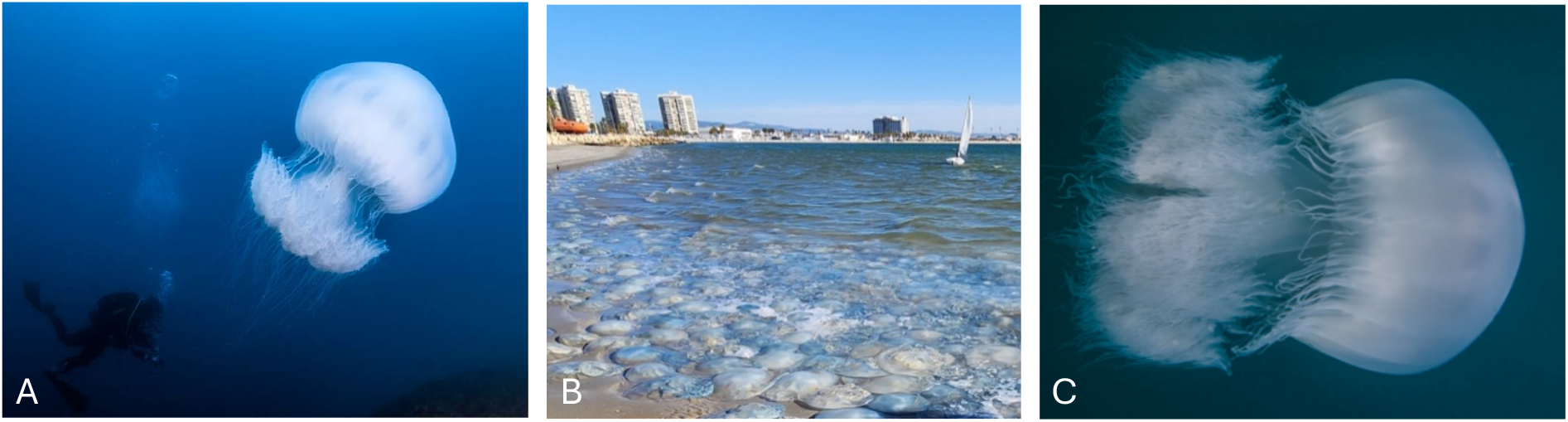
(A) *R. nomadica* in Israeli waters (photo: B.S. Rothman), (B) Stranded swarms of *R. nomadica* in Israel (photo: Z. Kuplik) (C) *R. nomadica* in Tanzanian waters, Mafia Island (photo: https://seaunseen.com/).

**Fig. 2.**
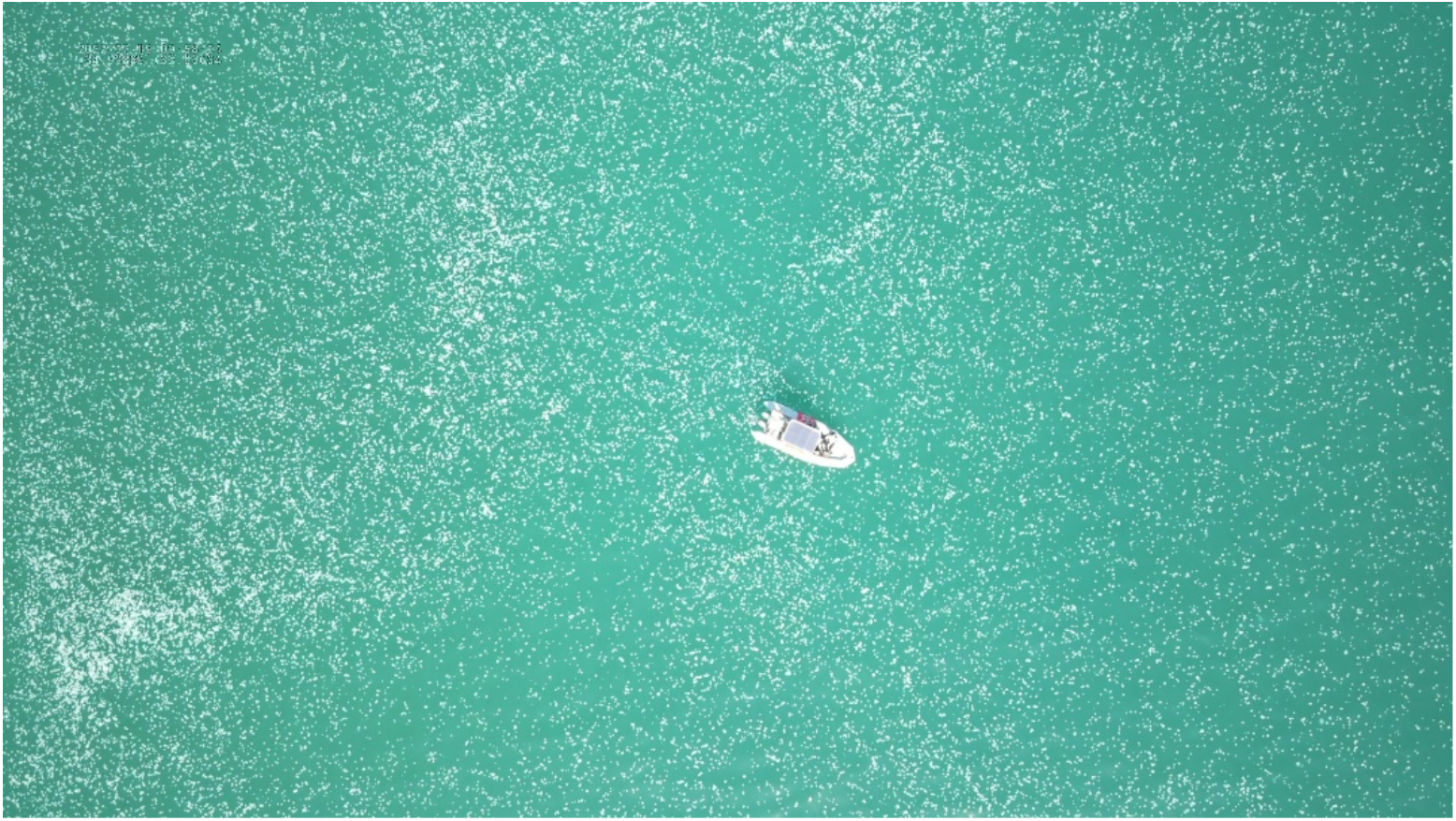
*R. nomadica* swarming off the coast of Haifa, Israel, July 2022 (photo: Rotem Sadeh)

*Rhopilema nomadica* is only one of many species that have been introduced to the Mediterranean; since opening in 1869, the Suez Canal has served as a corridor for introducing > 400 multicellular species (Galil et al. 2021). However, there are gaps in our understanding of its ecology, origin, and geographical expansion. Until recently, as there were no reports of *R. nomadica* in the Mediterranean Sea prior to the 1970s (Galil et al. 1990), it was assumed to have arrived around this time. However, a photograph taken in 1938 (Rakotsh, Fig. 3), shows a dense aggregation of *R. nomadica* in Haifa Port, Israel. Furthermore, despite it being considered a Lessepsian migrant, based on a specimen collected off Kamaran, Yemen, southern Red Sea, in 1939 (holotype, RMNH 7038) (Galil et al. 1990), only Berggren (1994) and Tahera & Kazmi (2015) reported *R. nomadica* outside the Mediterranean, and the report by Tahera & Kazmi (2015) was found to be a misidentification (Morandini & Gul 2016). Divers and researchers in eastern Africa have recently reported the presence of “nomad medusae” with similar morphology to *R. nomadica* near Dar es Salaam, Tanzania (Fig. 1C) (e.g. https://seaunseen.com/; Richmond 2002), suggesting this is the source of the Mediterranean population. However, this conjecture has yet to be tested.

**Fig. 3.**
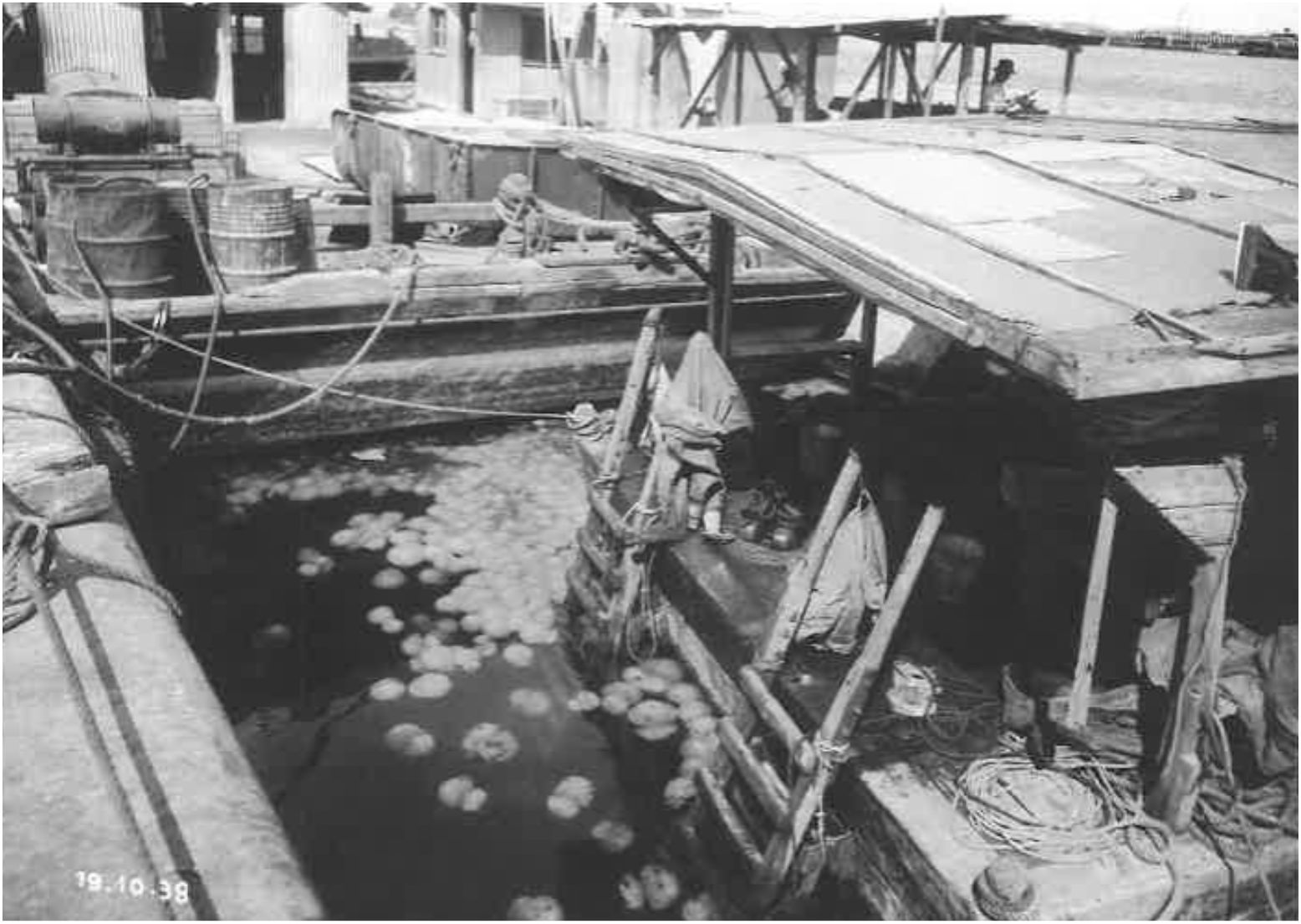
Yitzhak Rakotsh, Jellyfish in the Port. 19.10.1938. Silver print, 16.5 × 23 cm. From: Home Port, the story of Haifa Port, October 2002. Haifa Museums, Haifa City Museum. Courtesy Haifa Port. The species was identified by us and by André C. Morandini, an expert in scyphomedusa taxonomy, University of São Paulo, Brazil.

Here, we provide evidence that the “nomad medusa”, from the western Indian Ocean, is *R. nomadica*. We then explore why it has only relatively recently established persistent annual swarms in the Mediterranean. We propose that *R. nomadica* medusae were introduced into the Mediterranean over the last century, possibly soon after the Suez Canal was constructed. Moreover, we suggest that its apparent appearance in the late 1970s was due to changing environmental conditions, favoring the success of its reproductive benthic stage **[Box 1]**.

### Box 1.

*Rhopilema nomadica* has a metagenic life cycle composed of the free-swimming, planktonic medusa, and the benthic sessile polyp. Sexual reproduction occurs at the medusa stage and asexual reproduction occurs at the polyp stage. The medusa is gonochoric, releasing sperm or oocytes. Fertilization occurs in the water column. Fertilized eggs develop into free-swimming ciliated planula larvae that seeks a suitable surface to settle on. After settlement, the planula metamorphoses into a sessile polyp called ‘scyphistoma’. The scyphistoma then begins to reproduce asexually through podocyst formation, with the potential to create dense populations of scyphistomae within weeks. Given suitable conditions (e.g., temperature, food), the scyphistoma starts to produce medusae through a process called ‘strobilation’. The success of the benthic/scyphistoma stage is undoubtedly responsible for the formation of the medusa swarms (Lucas et al. 2012, Fu et al. 2014). Nevertheless, for most species this stage is cryptic, and scyphistoma populations have rarely been observed in the wild.

**Fig. 4.**
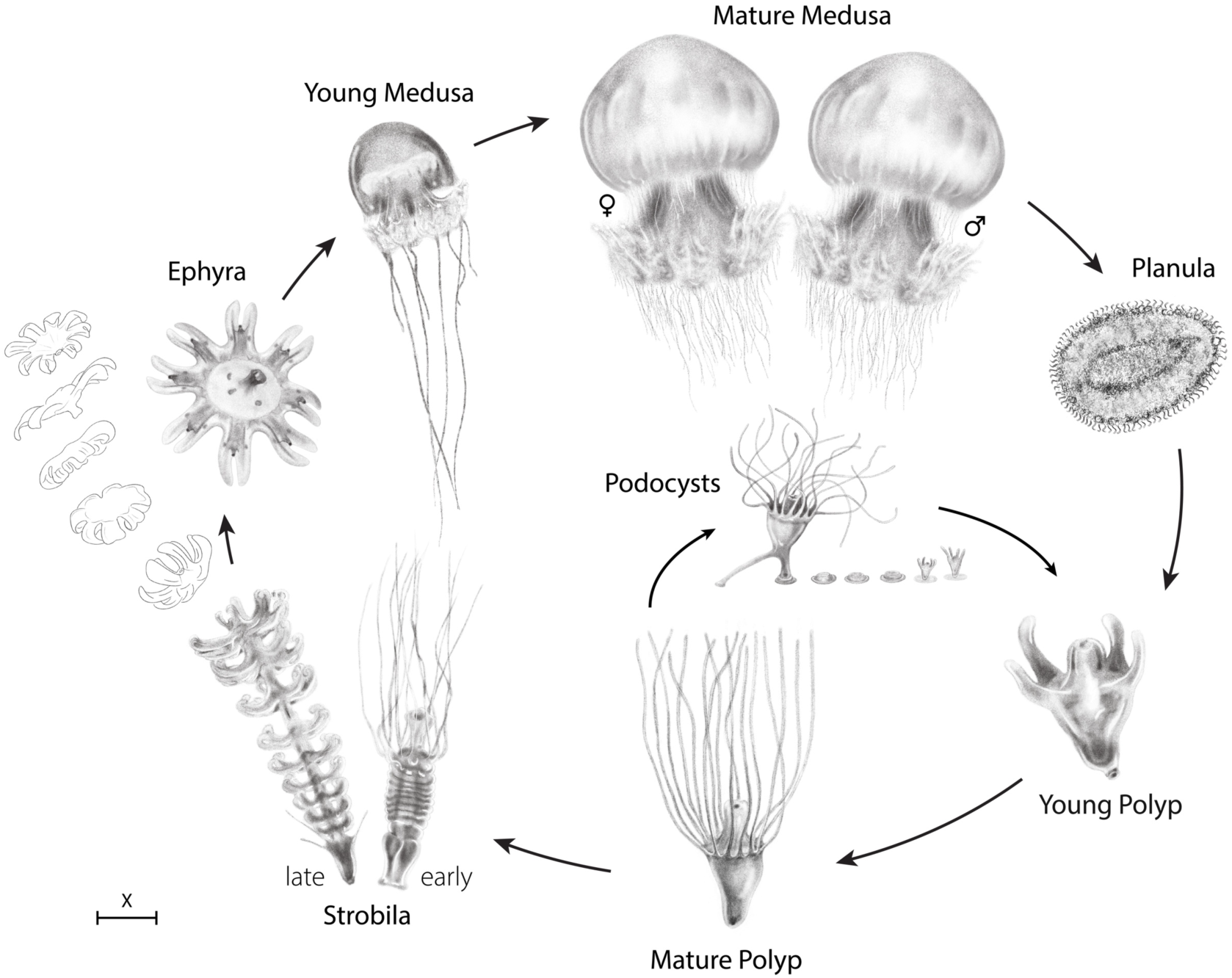
*Rhopilema nomadica* life cycle (clockwise). Rhopilema nomadica life cycle (clockwise). Scale for each of the life stages: Mature medusa X=10 cm, planula X=90 μm, young polyp X=100 μm, mature polyp X=700 μm, podocysts X=600 μm, strobila X=630 μm, ephyra X=800 μm, young medusa X=1.5 cm (Sketch by: Rahel Wachs)

### DNA sampling, amplification, and phylogenetic analyses

In order to understand the phylogenetic position of *R. nomadica* from Tanzania (three medusae stranded on the beach in Dar es Salaam, Tanzania 6°44’28.76”S, 39°16’33.39”E), a genus level phylogeny was reconstructed. For the molecular analyses, 217 individuals of *Rhopilema* were included, of which three were the newly collected specimens from Tanzania (labelled RN1P, RN2P, and RN3P) and 214 available specimens were retrieved from GenBank (48 of *R. esculentum*, 50 of *R. hispidum*, three of *R. verrilli*, and 113 of *R. nomadica*). It is noteworthy that all of the available sequences of *R. nomadica* were from the Mediterranean Sea solely, i.e. the newly generated Tanzanian sequences were the only ones from outside the Mediterranean region. Additionally, four sequences of the genus *Rhizostoma* (also from the family Rhizostomatidae) were retrieved from GenBank to root the phylogenetic trees (two sequences each for *R. luteum* and *R. pulmo*).

DNA was extracted using a CTAB-phenol/chloroform protocol (Dawson & Jacobs, 2001). The samples were PCR-amplified and bi-directionally sequenced for the mitochondrial protein-coding gene fragment of cytochrome oxidase subunit 1 (COI). The mitochondrial COI fragment was amplified using both the species-specific primer set RNF RNR and the wide range jellyfish primer set LCOjf HCOcato (Table 1). Both strands of the PCR products were sequenced, and chromatographs were checked, assembled, and edited using Geneious v.7.1.9 (Biomatter Ltd.). The Mafft plugin of Geneious was used with default settings to align all sequences. The sequences were translated to amino acids and no stop codons were detected.

**Table 1.**
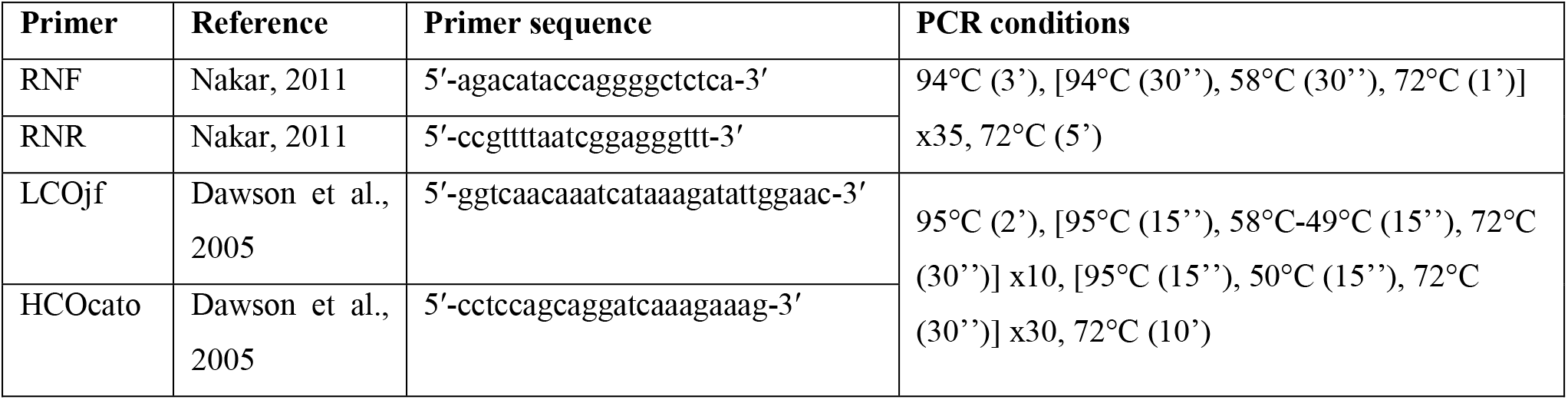
Data on the primers used in this study, including their orientation, sequences, references and PCR conditions.

For the phylogenetic analyses, the COI dataset of 655 bp was analysed under Maximum Likelihood (ML) and Bayesian Inference (BI) frameworks. The ML analyses were performed using IQ-TREE v. 1.6 (Nguyen et al., 2015) through the web interface (Trifinopoulos et al., 2016). The best-fit substitution model was selected automatically for each partition during the analysis. Branch support was assessed with the Shimodaira-Hasegawa-like approximate likelihood ratio test (SH-aLRT; Guindon et al., 2010) and the ultrafast bootstrap (UFBoot; Hoang et al., 2018), both with 1,000 replicates. The BI analyses were carried out using MrBayes v.3.2.7 (Ronquist et al., 2012). The best substitution model was determined using JModelTest v.2.1.7 (Darriba et al., 2012; Guindon & Gascuel, 2003) which resulted in the GTR+G model. Nucleotide substitution model parameters were unlinked across partitions and the different partitions were allowed to evolve at different rates. Two simultaneous parallel runs were performed with four chains per run (three heated, one cold) for 10 million generations with sampling frequency of every 1,000 generations. Stationarity was determined by the standard deviations of the split frequencies being lower than 0.01. The standard deviation of the split frequencies between the two runs and the Potential Scale Reduction Factor (PSRF) diagnostic were examined. The first 25% of trees were conservatively discarded as burn-in. Nodes with SH-aLRT ≥ 80, UFBoot ≥ 95, and a Bayesian posterior probability ≥ 0.95 were considered strongly supported. We calculated inter- and intraspecific uncorrected *p*-distance between *Rhopilema* and *Rhizostoma* species for the COI mitochondrial fragment, with pairwise deletion, in MEGA11 (Tamura et al., 2021).

## Results and Discussion

The results of the phylogenetic analyses, based on the COI dataset and using BI and ML analyses, produced similar trees differing mostly in the less supported nodes at the intraspecific level (Figure 5). *Rhopilema* was recovered as monophyletic (SH-aLRT = 97.7/ UFBoot = 100/ Bayesian posterior probability = 1.0; support values are given in the same order hereafter). Monophyly of all species of the genus was strongly supported (*R. esculentum*: 100/100/1.0; *R. verrilli*: 99.2/100/1.0; *R. hispidum*: 99.9/100/1.0; *R. nomadica*: 100/100/1.0). The species *R. esculentum* was recovered as the sister taxon to all the other *Rhopilema* species included in the analysis, and *R. verrilli* was the second species to branch off the tree (91.6/98/1). A sister taxa relationship between *R. hispidum* and *R. nomadica* was recovered with relatively weak support (58.9/85/0.95). The species *R. nomadica* showed shallow levels of genetic differentiation and the three Tanzanian specimens were all phylogenetically situated within the species, which is represented by specimens from the Mediterranean Sea. Among these three Tanzanian specimens, two (RN1P and RN3P) had identical sequences, whereas the third (RN2P) is differentiated from both at only one site, at position 307.

**Fig. 5.**
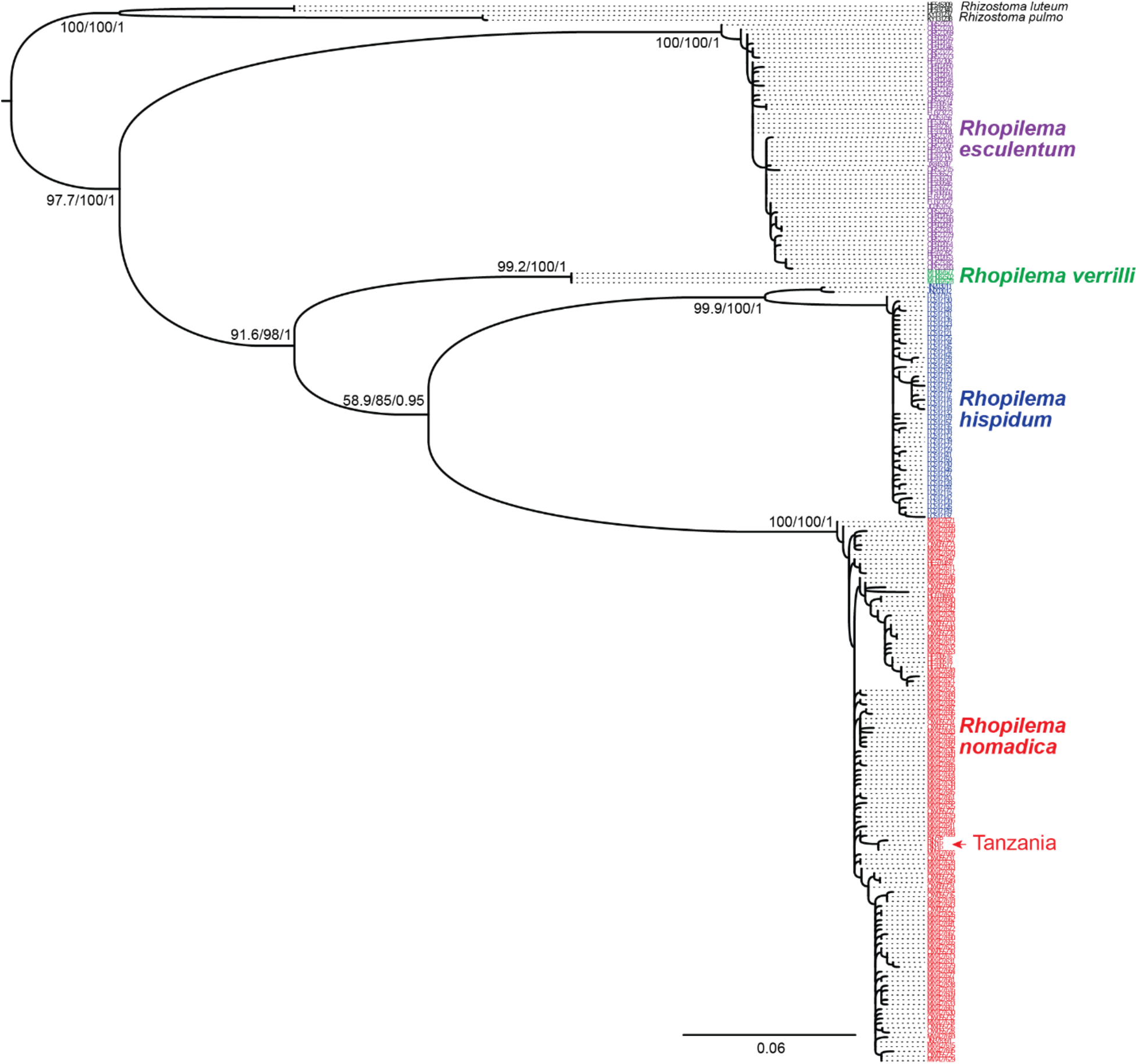
Phylogenetic position of the Tanzanian specimens of *R. nomadica*. The phylogenetic tree was derived from the Maximum likelihood analysis of *Rhopilema* (each species is highlighted in color: *R. esculentum* in purple; *R. verrilli* in green; *R. hispidum* in blue; *R. nomadica* in red) and *Rhizostoma* outgroup taxa. Colored arrow indicates the location of the three Tanzanian specimens of *R. nomadica*. Numbers near the nodes indicate support values in the following order: SH-aLRT, UFBoot, posterior probability resulting from the Bayesian analysis.

The uncorrected genetic distance in the COI fragment ranged between 15–17.4% among *Rhopilema* species, and 15–18.4% between *Rhopilema* and *Rhizostoma*. The lowest distance among *Rhopilema* species was 15% between *R. verrilli* and *R. hispidum* and the highest was 17.4% between *R. verrilli* and *R. esculentum*. The level of genetic variability within *Rhopilema* species was relatively low and ranged between 0–3% (*R. esculentum*, 3%; *R. nomadica*, 1%; *R. hispidum*, 1%; *R. verrilli*, 0%).

The phylogenetic analysis confirms that the medusae collected from Dar es Salaam, Tanzania, were *R. nomadica* (Fig. 5). This is the first genetic indication that the *R. nomadica* in the Mediterranean originated in the Indo-Pacific region. We suggest that, like many other introduced species, *R. nomadica* arrived in the eastern Mediterranean via ballast water in ships from east African waters.

Galil et al. (1990) reported that *R. nomadica* was found in the Red Sea and that it entered the Mediterranean via the Suez Canal, yet there have been no published reports of its presence elsewhere in the Red Sea, nor anywhere else, aside from the Mediterranean. It was only in 1994, in the waters of Mozambique, 2500 km south of Tanzania, that a medusa, serving as a host to a pelagic shrimp, was identified as *R. nomadica* (Berggren 1994); however, this identification was based on photographs (unpublished), taken while snorkeling (Berggren, personal communication). Hence, our results are the first to provide sound molecular evidence for the presence of *R. nomadica* in the waters of East Africa. It appears that this medusa, now identified as *R. nomadica*, is seasonally abundant offshore Dar es Salaam, Tanzania (A. Sutton, personal communication), generally between October and December.

In common with other ephemeral jellyfish that have a metagenic life cycle, *R. nomadica* may exist and persist as cryptic populations (albeit in small numbers) in a variety of habitats; and may remain unnoticed if they do not swarm or when no one is looking for them. In 2000, a swarm of the previously unreported rhizostome *Phyllorhiza punctata*, occurred in the northern Gulf of Mexico (Graham et al. 2003). This scyphomedusa likely entered the Atlantic Ocean from the Pacific through the Panama Canal some 50 years earlier (Graham et al. 2003), but the cryptic population was undetected until it bloomed. Likewise, in Brazil, the first report of the upside-down jellyfish (*Cassiopea*) was published in 2002 (Migotto et al. 2002); however, based on molecular findings, *Cassiopea andromeda* may have existed unnoticed in Brazilian waters for as long as 500 years before the genus was first described in Brazil (Morandini et al. 2017).

It seems that the date of first observation of a migrant species does not necessarily correspond to the date of introduction as it may take some time for this species to be detected (Walsh et al. 2016). This applies even more strongly to scyphozoans, which have a metagenic life cycle. As Walsh et al. (2016) suggested, a long-time lag between introduction and detection, as observed for *R. nomadica*, is not uncommon, but the real challenge is to identify the reasons for the proliferation of a species that had previously maintained a low population density.

Here we propose that a cryptic population of *R. nomadica* was found in the Mediterranean years before it was described, and it became apparent only after changes in environmental conditions. This hypothesis is supported by an observation in 2014, when a sudden bloom of *C. andromeda* occurred in our flowing seawater aquaria. Although it has been documented as one of the first Lessepsian migrants (Maas 1903), reports of the medusa stage of *C. andromeda* in the Mediterranean coastal waters of Israel are very rare. A short search in our seawater aquaria revealed a rock, covered with *C. andromeda* polyps that was collected at sea and left in one of the tanks. Most likely, the protected waters, and possibly abundant food from aquaria feedings, led to strobilation and to the outbreak of this species. Following this introduction, *C. andromeda* became a dominant ‘fouling’ organism in the flowing seawater system, where polyps proliferate and strobilate, and ephyrae are released seasonally.

Using *C. andromeda*’s proliferation in the laboratory as a case study, we propose a similar scenario for *R. nomadica* in the wild. We know what conditions *C. andromeda* needs to bloom, but what conditions does *R. nomadica* need? Perhaps therein lies the answer to the apparent absence of *R. nomadica* medusae until the 1970s. The findings of Edelist et al. (2022) concur with our assumption that the origin of *R. nomadica* swarms in the eastern Mediterranean is in the Nile Delta, suggesting that this is the region where many of the polyps reside and release ephyrae. Success of the polyp stage is often inhibited by non-optimal temperatures, insufficient food quality and quantity, or a combination of these (Lucas et al. 2012, Chi et al. 2019, Loveridge et al. 2021, Dror & Angel 2024). Therefore, we searched for environmental changes, on a regional scale that could have affected polyp reproduction and the subsequent medusa populations.

The opening of the Aswan High Dam in 1965 caused a change in the nutrient regime in eastern Mediterranean waters (Halim et al. 1976), which may have led to changes in *R. nomadica* abundances. Prior to 1965, nutrient enrichment from the Nile led to Mediterranean phytoplankton blooms, most notably the August-October flood (Sharaf 1977). Unlike other parts of the oligotrophic eastern Mediterranean (Azov 1991), these plankton-rich Egyptian coastal waters, dominated by diatoms (>90%) (Halim 1960, Aleem & Dowidar 1967, Dowidar 1984), were the source of rich planktivorous fisheries. Shortly after the High Dam became operational (early 1970s), the phytoplankton composition changed to a nearly balanced mix of both diatoms and small flagellates (Dowidar 1984), followed by the collapse of the sardine population (Aleem 1972). Even more than a decade later, in the early 1980s, it was found that in the south-eastern Mediterranean, 60-80% of chlorophyll-a biomass consisted of small (nano- and pico-) phytoplankton (Dowidar 1984), such as flagellates, in contrast with the large diatoms of pre-Aswan High Dam construction. These changes in the lower trophic level may have led to an increase in ciliates (Stoecker & Capuzzo 1990) that feed on small flagellates (Jonsson 1986) and are in turn grazed by zooplankton (Nixon 2003). Ciliates may also sustain benthic scyphozoan polyps and ephyrae (Kamiyama 2011, 2013, 2018), enabling a substantial increase in their abundances and subsequently to adult jellyfish blooms, as observed in the late 1970s and early 1980s (Galil et al. 1990). Another factor to be considered is the ongoing warming trend of the Mediterranean Sea (Pastor et al. 2020). Recently, Dror and Angel (2023) showed that the rising sea surface temperatures are a major factor in polyp life history, positively affecting both polyp survival and asexual reproduction. This, coupled with a favorable diet, could explain the noticeable change in the pelagic population of *R. nomadica*, leading to the massive annual swarms that now characterize the eastern Mediterranean.

The new data we provide here support the initial assumption that this species was introduced to the Mediterranean from the Indian Ocean, though it is not clear whether this was via swimming through the Suez Canal or via ballast water. Both the route of introduction/invasion and the dynamics of its proliferation need to be further investigated, via polyp feeding trials, prey preferences and growth trials, and population genetics to better understand the relationship between eastern African *R. nomadica* populations and those of the Mediterranean. Improved understanding of these could provide us with essential information to help track its population dynamics and predict its future distribution to new habitats. The fact that *R. nomadica* became a dominant species in the eastern Mediterranean only in the late 1970s, long after it was first documented in 1938, raises interesting questions concerning the ecological dynamics involved. Nevertheless, we may never know if it was the opening of the Aswan High Dam that triggered a series of ecological changes that resulted in *R. nomadica* becoming the dominant species in the Levant.

## Acknowledgements

We are hugely grateful to Prof. David Montagnes for his suggestions on the draft manuscript. We thank Rafael Yavetz, Liel Uziyahu, and the Mevo’ot-Yam high-school for the use of their flowing seawater facilities, their support and their invaluable technical assistance. We are grateful for the technical support of Ms. Victoria Fidel of Dr. Omri Bronstein’s lab, the Steinhardt Museum of Natural History, Tel Aviv University, with the DNA extraction.

